# Ongoing Production of Tissue-Resident Macrophages from Hematopoietic Stem Cells in Healthy Adult Macaques

**DOI:** 10.1101/2022.12.19.521067

**Authors:** Andrew R. Rahmberg, Chuanfeng Wu, Taehoon Shin, So Gun Hong, Luxin Pei, Heather D. Hickman, Cynthia E. Dunbar, Jason M. Brenchley

**Affiliations:** Barrier Immunity Section, Lab of Viral Diseases, Division of Intramural Research, National Institute of Allergy and Infectious Diseases, NIH, Bethesda, MD; Translational Stem Cell Biology Branch, National Heart, Lung, and Blood Institute, National Institutes of Health, Bethesda, MD; Viral Immunity and Pathogenesis Unit, Lab of Clinical Immunology and Microbiology, NIAID, NIH, Bethesda, MD

## Abstract

Macrophages are critical orchestrators of tissue immunity, from the initiation and resolution of antimicrobial immune responses to the subsequent repair of damaged tissue. Murine studies have demonstrated that tissue-resident macrophages are comprised of a mixture of yolk sac-derived cells that populate the tissue before birth and hematopoietic-derived replacements that are recruited in adult tissues both at steady-state and in increased numbers in response to tissue damage or infection. While resident macrophages in some murine tissues are readily turned over and replaced, other tissues primarily retain their original, yolk sac-derived complement of macrophages. How this translates to species that are constantly under immunologic challenge, such as humans, is unknown. To understand the ontogeny and longevity of tissue-resident macrophages in nonhuman primates (NHPs), we employ a model of autologous HSPC transplantation with HSPCs genetically modified to express individual bar-coded markers allowing for subsequent analysis of clonal differentiation of leukocyte subsets. We study the contribution of HSPC to tissue-macrophages and their clonotypic profiles relative to leukocyte subsets across tissues and peripheral blood. We also use in vivo bromodeoxyuracil infusions to monitor tissue macrophage turnover in NHPs at steady state. We find that, in all anatomic sites we studied, HSPC contribute to tissue-resident macrophage populations. Their clonotypic profile is dynamic and overlaps significantly with the clonal hierarchy of contemporaneous monocytes in peripheral blood. Moreover, we find evidence of tissue-macrophage turnover at steady state in otherwise unmanipulated NHPs. These data demonstrate the life span of tissue-resident macrophages can be limited and they can be replenished from HSPCs. Thus, in primates not all yolk-sac derived tissue-macrophages survive for the duration of the host’s life.

## Introduction

Macrophages are diverse and functionally important. Evolutionarily conserved across most species of the Chordata phylum, macrophages are one of the “oldest” leukocyte subsets and have the highest degree of plasticity (Jenkins et al., 2011). Macrophages play supportive roles in multiple aspects of physiology including infections and tissue repair. Their function and phenotype depend upon anatomical location as well as response to infection, tissue damage, adjacent tumor cells, or other environmental stressors, based on both intrinsic factors such as ontogeny and degree of differentiation as well as extracellular cues.

Tissue macrophages are important targets for many RNA and DNA viruses (reviewed in (Estes et al., 2018)) and are hypothesized to be long-lived reservoirs for certain viruses such as human immunodeficiency virus (HIV) and simian immunodeficiency virus (SIV) (DiNapoli et al., 2016). Moreover, tissue-resident macrophages infected with some viruses can resist immunological killing by NK cells and CD8 T cells (Clayton et al., 2018; Clayton et al., 2021). Design of strategies aimed at completely depleting reservoirs of these pathogenic viruses depends on the life span and ontogeny of infected macrophages and their ability to be cleared by the immune system.

Murine macrophages first arise during embryonic development in the yolk sac, then populate the entire embryo once circulation is established and persist into adulthood, as demonstrated via sophisticated fate-mapping studies (Ginhoux et al., 2010; Gomez Perdiguero et al., 2015). Macrophages derived from the yolk sac or fetal liver reside in organs such as the brain (as microglia cells), pancreas, spleen, liver (as Kuppfer cells), and kidney are thought to be very long-lived, perhaps for the life span of the host (Ginhoux *et al*., 2010). Macrophages can also differentiate *in vitro* from hematopoietic stem and progenitor cell (HSPC)-derived circulating monocytes, and particularly under stress, there is evidence they can replace or replenish tissue-resident macrophages (Elmore et al., 2014; Sailor et al., 2022; Stutchfield et al., 2015; Theurl et al., 2016). Once in tissues, recent murine studies suggest that macrophages can divide to maintain homeostasis (Jenkins *et al*., 2011; Saijo and Glass, 2011; Schulz et al., 2012). In specific-pathogen-free mice, the extent and timing of monocyte replacement varies dramatically by tissue, murine strain, and conditions of the experiment, with some tissues, such as the brain, having almost no turnover (Mildner et al., 2007). By contrast, the dermis is thought to be primarily populated by monocyte-derived macrophages (Scott et al., 2014; Tamoutounour et al., 2013). Thus, the unique balance of long-lived versus newly differentiated macrophages has the potential to profoundly impact pathogen handling at the tissue level. Viral infection of lymph node macrophages leads to their rapid death, as well as the death of uninfected macrophage neighbors (Gaya et al., 2015). These macrophages are rapidly replaced by incoming monocytes (Mondor et al., 2019). Thus, there exists a continuum of scenarios for macrophage handling of pathogens that centers around their longevity and understanding their turnover dynamics of particular interest and importance.

Emerging data suggest that environmental variables can dramatically influence the immunological development and responses of the host. Inbred mouse strains maintained in captivity are immunologically disparate from wild mice and respond to pathogenic infections very differently (Beura et al., 2016; Rosshart et al., 2017). Indeed, the interplay between endogenous host traits, their microbiome, other environmental factors, and infection with pathogens represents an area of active research (Ansaldo et al., 2021). Macrophage development and replacement remain to be explored in wild or “dirty” mice. Further, it is unclear how macrophage turnover dynamics in laboratory mice would translate to longer-lived, pathogen-exposed species such as primates.

To explore the ontogeny and longevity of tissue macrophages in outbred populations of NHPs with high relevance to human biology and disease based on life span and characteristics of both HSPC and immune cells, we performed autologous HSPC transplantation after genetically modifying the HSPCs with a lentiviral vector expressing green fluorescent protein and a bar-coded genetic marker (Cordes et al., 2021; Koelle et al., 2017; Wu et al., 2014). As the HSPCs were transduced with lentiviral vectors containing a library of individual bar-coded genetic markers, this approach allowed us to explore the clonal composition of individual leukocyte subsets, including tissue-resident macrophages, in multiple anatomic sites, relative to their clonal composition in peripheral blood (Koelle *et al*., 2017; Wu *et al*., 2014). To assess turnover of tissue-macrophages in animals that did not receive the stressor of transplantation conditioning, we infused bromodeoxyuracil (BrdU) into healthy animals and followed the emergence and loss of BrdU^+^ tissue-resident macrophages longitudinally.

## Material and Methods

Fifteen adult rhesus macaques (*Macaca mulatta)* were studied, including four that underwent autologous barcoded hematopoietic stem progenitor cells (HSPCs) transplantation. This study was carried out in strict accordance with the recommendations described in the Guide for the Care and Use of Laboratory Animals of the National Institutes of Health, the Office of Animal Welfare, and the U.S. Department of Agriculture (Council, 2011). All animal work was approved by the NIAID and NHLBI Division of Intramural Research Animal Care and Use Committees in Bethesda, MD (protocols H-0136 and LVD-26E). The animal facilities are accredited by the American Association for Accreditation of Laboratory Animal Care. All procedures were carried out under ketamine anesthesia by trained personnel under the supervision of veterinary staff, and all efforts were made to maximize animal welfare and to minimize animal suffering in accordance with the recommendations of the Weatherall report on the use of nonhuman primates (Weatherall, 2006). Animals were housed in adjoining individual or paired primate cages, allowing social interactions, under controlled conditions of humidity, temperature, and light (12-hour light/12-hour dark cycles). Food and water were available *ad libitum*. Animals were monitored twice daily and fed commercial monkey chow, treats, and fruit twice daily by trained personnel. Environmental enrichment was provided in the form of primate puzzle feeders, mirrors, and other appropriate toys.

### Autologous barcoded HSPC transplantation

Barcoded lentivirus libraries were generated and used for rhesus macaque CD34^+^ HSPCs transduction as previously described, with library diversity confirmed to be sufficient to ensure a >95% chance that each barcode would be present in no more than one engrafting HSPC (Lu et al., 2011; Wu *et al*., 2014). Autologous transplantation with lentivirally-transduced CD34^+^ HSPCs was performed as previously described (Donahue et al., 2005; Wu *et al*., 2014). Briefly, rhesus macaque HSPCs were mobilized from the bone marrow by treatment with granulocyte colony-stimulating factor (G-CSF) 10-15 mg/kg/day (Amgen, Thousand Oaks, CA) for 5 days and AMD3100 (plerixafor) 1 mg/kg (Sigma, St. Louis, MO) on the morning of the fifth day. Collection of peripheral blood mononuclear cells (PBMCs) via apheresis was performed followed by immunoselection of CD34^+^ cells. CD34^+^ cells were cultured overnight on RetroNectin-coated plates (Takara, T100B, Mountain View, CA) in X-VIVO 10 media (Lonza, Rockland, ME) supplemented with 100 ng/mL of recombinant human flt3 ligand (Miltenyi Biotec, Auburn, CA), stem cell factor (SCF, Miltenyi Biotec), and thrombopoietin (TPO, Miltenyi Biotec). The next day concentrated barcoded lentiviral vector was added at a multiplicity of infection of 25 in the presence of 4 μg/mL protamine sulfate (Sigma). 24 hr following addition of the vector library, cells were collected and reinfused intravenously into the autologous rhesus macaque. Each animal received total-body irradiation at a dose of 500 cGy/day for 2 days prior to the day of cell reinfusion as pre-transplantation conditioning.

### Tissue processing and cell purification

Cells were isolated from peripheral blood by density gradient centrifugation over lymphocyte separation media (MP Biomedicals, #0850494-CF), washed in PBS and counted. Four core biopsies each of spleen and liver were obtained during laparotomy, and ten pinch biopsies from colon and jejunum were obtained during endoscopy. Tissue biopsies were homogenized into single cell suspensions as previously reported (Ortiz et al., 2018). Bronchioalveolar lavage was performed by instilling 150ml of saline into the lung followed by collection via suction. Cells were stained with amine-reactive viability dye (Thermo Fisher Scientific, #L34966) and a panel of antibodies (Table S1) for 20 minutes at 4°C. Cells were washed in PBS and fixed with 1% paraformaldahyde prior to analysis or sorting. Cells were sorted on a FACSymphony S6 machine (Becton-Dickinson).

### Confocal microscopy

Sorted cells were resuspended in PBS and plated in chambered coverglasses (LabTek). Images of individual cells were acquired at high magnification using a 63X objection and a Leica SP8 inverted confocal microscope equipped with HyD-detectors (Leica Microsystem). A total of 30 images were acquired per cell type. Final image processing was done using Imaris (Bitplane).

### Barcode retrieval

Genomic DNA from sorted cell populations of greater than 200,000 cells was extracted using the DNeasy Kit (Qiagen). DNA from cell populations less than 200,000 cells was processed via direct lysis using DirectPCR lysis buffer (Viagen Biotech). 200ng DNA or the whole cell lysis from each cell population underwent 28-cycle PCR using Phusion High-Fidelity DNA Polymerase (ThermoFisher) as described (Wu et al., 2018; Wu et al., 2014). A universal reverse primer and a set of forward primers with unique i5 indexes were used to allow multiplexing for sequencing. Following gel-purification, samples were pooled and sequenced on an Illumina HiSeq 2500, HiSeq 3000 or NovaSeq 6000 system (Wu et al., 2018b; Wu *et al*., 2014). Sequencing output was processed using custom Python and R code to quantitate individual barcode frequency (available at: www.github.com/dunbarlabNIH/ and described in (Espinoza et al., 2021)), validated to accurately quantitate fractional contributions of each clone within highly polyclonal cellular populations (Koelle *et al*., 2017; Wu *et al*., 2018b; Wu *et al*., 2014).

### BrdU infusion and analysis

BrdU (Sigma) was prepared at a concentration of 10 mg/mL in Hank’s balanced salt solution (Sigma), sterile filtered, and administered intravenously to rhesus macaques at 30mg/kg/day for 5 consecutive days. Blood was drawn prior to infusion and on days 7, 9, 10, 11, and 24 after the start of infusions. Biopsies of spleen, inguinal lymph node, and mesenteric lymph node were collected on days 10 and 24 following the initial infusion. Cells were analyzed for BrdU positivity by flow cytometric analysis using a Becton Dickinson Fortessa.

### Statistical analyses

Paired t tests were used to determine statistical significance (Prism). Spearman correlations were used to determine relationships between variables and 95% confidence intervals were used to determine clustering in principal coordinate plots. Correlation heatmaps were produced via our BarcodetrackR custom software (Espinoza *et al*., 2021).

## Results

### Marking frequencies of HSC-derived leukocytes in tissues

We performed autologous transplantation of rhesus macaques with CD34^+^ HSPCs transduced with lentiviral vectors expressing copepod green fluorescent protein (GFP) and containing a high diversity library of genetic barcodes (Cordes *et al*., 2021; Wu *et al*., 2014)(Figure 1A). This vector design allows utilization of GFP as a marker gene demonstrating derivation of various cell lineages in tissues from transplanted HSPCs, as well as lineage tracing of HSPC output at a clonal level via quantitative barcode retrieval. Transplantation parameters, including HSPC cell doses, transduction efficiency assessed in infused CD34^+^ HSPC, and tissue collection time points post-transplantation are summarized in Table 1. Analyses of peripheral blood (PB) and bone marrow (BM) longitudinal clonal dynamics in the four transplanted macaques included in this study have been in part previously published (Fan et al., 2020; Koelle *et al*., 2017; Wu *et al*., 2018b; Wu *et al*., 2014).

**Table 1:** Animal transplantation parameters and sample collection timing.

|  | **JM82** | **ZK22** | **ZJ31** | **ZG66** |
| --- | --- | --- | --- | --- |
| CD34+ transplanted dose (millions) | 91 | 82 | 23 | 48 |
| CD34+ transplanted dose/kg (millions) | 7.2 | 7.2 | 4.1 | 8.5 |
| % GFP+ infused CD34+cells | 34% | 31% | 35% | 35% |
| Infused GFP+ cells (millions) | 30.6 | 25.2 | 8.0 | 16.7 |
| Sample timepoints included (months post-transplantation) | 1, 12, 29, 46, 47, 48, 49 | 2, 13, 30, 45, 62, 63 | 2, 35, 63, 74, 75, 76 | 1, 36, 76, 104, 106 |

**Figure 1:**
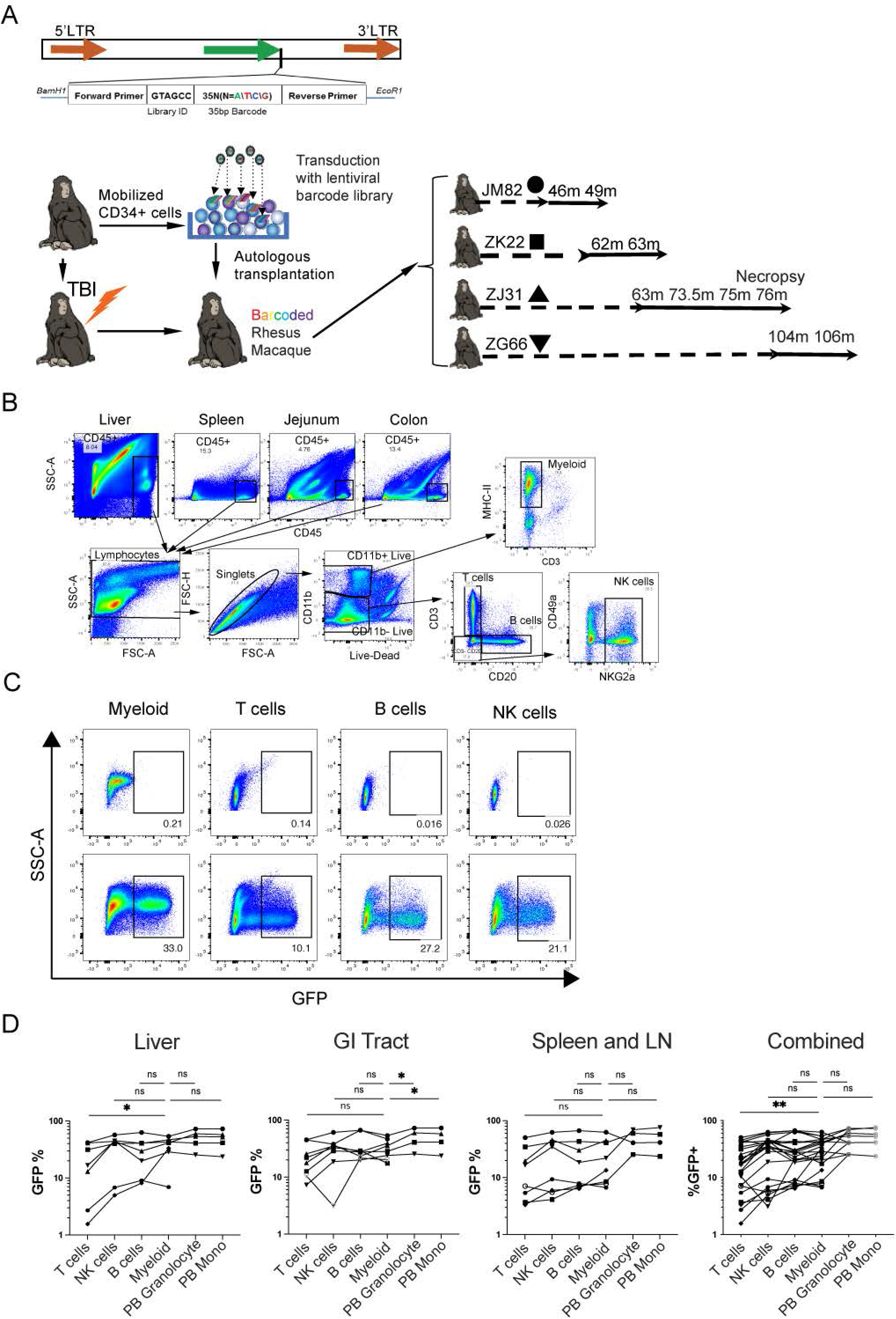
Rhesus macaque autologous barcoded HSPC transplantation and detection of GFP+ cells in tissues. **(A)** Rhesus macaque HSPCs were transduced with a barcoded lentiviral library and transplanted into the autologous macaque following conditioning with 1000 rads total body irradiation. Blood and tissue were sampled periodically to assess the clonal composition of HSPC-derived cells. (**B)** Representative flow cytometric gating strategy to measure GFP expression among leukocyte subsets. (**C)** The expression of GFP was determined by flow cytometry in tissue mononuclear cell suspensions following gating for CD45 positive cells, lymphocytes, and then lineage-defining markers. Representative liver samples from an untreated control animal (top) and barcoded animal ZJ31 sample collected 76 months post-transplantation (bottom). (**D)** The percent GFP positive hematopoietic cells in liver, gastrointestinal tract, or lymphoid tissues for the listed cell lineages is shown. Each individual animal is designated with a different symbol as depicted in A) and a line connects samples obtained at the same time point. For most tissue-resident myeloid cell %GFP positivity was not significantly different compared to B or NK cells in the same tissue or in granulocytes or monocytes in the peripheral blood, and was significantly higher than T cells in the tissue (2-way ANOVA with Dunnett’s multiple comparisons test, lines compare tissue-resident myeloid cells with other leukocyte subsets) * P<0.05, ** P<0.01.

We sampled tissue-resident leukocytes from multiple tissues including liver, spleen (via laparotomy), lymph node (via surgical excision), jejunum and colon (via endoscopy), lung (via broncheoalveolar lavage (BAL)), and PB at several time points post-transplantation to assess the degree to which individual myeloid and lymphoid lineage populations had been replaced by output from the transplanted, genetically marked HSPCs. Flow cytometry for GFP expression in mononuclear cell leukocyte populations, including T cells, B cells, NK cells and tissue myeloid cells (macrophages), was performed (representative gating strategy in Figure 1B and representative GFP expression in subsets shown in Figure 1C). Tissue macrophages were defined as CD45^+^CD11b^+^major histocompatibility class (MHC) II^+^ CD3^-^CD20^-^NKG2A^-^, T cells were defined as CD45^+^CD3^+^CD20^-^NKG2A^-^CD11b^-^, NK cells were defined as CD45^+^NKG2A^+^CD3^-^ CD11b^-^CD20^-^ and B cells were defined as CD45^+^CD20^+^CD11b^-^NKG2A^-^CD3^-^. Consistent with previous reports from our group and others, mature T cells in both PB and in most tissue were less frequently GFP^+^ and thus derived from transplanted transduced HSPCs compared to other leukocyte subsets (MacVittie et al., 2014; Wu *et al*., 2014) (Figure 1D), presumably due to decreased thymic output of new HSPC-derived T cells in adult macaques transplanted following total body irradiation conditioning and peripheral expansion of non-transplanted T cells surviving irradiation.

Notably, in all anatomic sites we were able to detect GFP^+^ tissue-resident macrophages (Figure 1C). Moreover, the frequency of GFP^+^ macrophages was as high as in any other population of tissue leukocytes and higher than the frequencies of GFP^+^ T cells (Figure 1D). Most importantly, the frequencies of GFP^+^ macrophages in tissues were no different than the frequency of GFP^+^ PB monocytes sampled concurrently (Figure 1D and data not shown). Indeed, tissue-resident macrophages were as frequently GFP^+^ as circulating neutrophils (Figure 1D), which have an estimated half-life of less than one week in humans. Thus, tissue-resident macrophages sampled years following transplantation were equally derived from transplanted HSPCs compared to other leukocyte subsets resident in liver, lymphoid tissues, mucosal tissues, or circulating in blood.

We have previously found that macrophage-mediated phagocytosis of other leukocyte subsets can confound analysis of tissue-resident macrophages (Calantone et al., 2014; DiNapoli et al., 2017). Thus, one potential explanation for GFP^+^ macrophages in tissue is phagocytosis of other GFP^+^ leukocytes, rather than derivation from HSPCs and migration into tissues post-transplantation. Phagocytosis of GFP^+^ leukocytes would be predicted to result in accumulation of GFP protein specifically within phagosomes and lysosomes, but not the cytoplasm. To understand the cellular organization of GFP within GFP^+^ macrophages relative to GFP^+^ T cells, we flow cytometrically sorted T cells and macrophages from the liver of one of our transplanted animals and analyzed GFP localization by confocal microscopy (Figure 2). We could find evidence of GFP localization within intracellular organelles of tissue-resident macrophages when focused on the organelles planes (Figure 2A), but we could also find clear evidence of dispersed cytoplasmic GFP (Figure 2B). Moreover, GFP localization patterns were similar within sorted T cells, i.e. both within intracellular organelles or dispersed across the cytoplasm) (Figure 2C). Moreover, myeloid cell phagocytosis of T cells results in the accumulation of rearranged TCR DNA in myeloid cells (Calantone *et al*., 2014). Thus, we measured levels of rearranged TCR DNA within isolated T cells and macrophages from liver and BAL of two of our transplanted animals. As expected, all sorted T cells had detectable rearranged TCR DNA (Figure 2D). In both macrophage samples we could find detectable, but very low, levels of rearranged TCR DNA in tissue-resident myeloid cells, more than two logs lower than present in T cells (Figure 2D). Taken together, it was clear that phagocytosis of other GFP^+^ leukocytes was not responsible for the majority of GFP positivity we detect within tissue-resident macrophages and instead supports their development from transplanted HSPCs.

**Figure 2.**
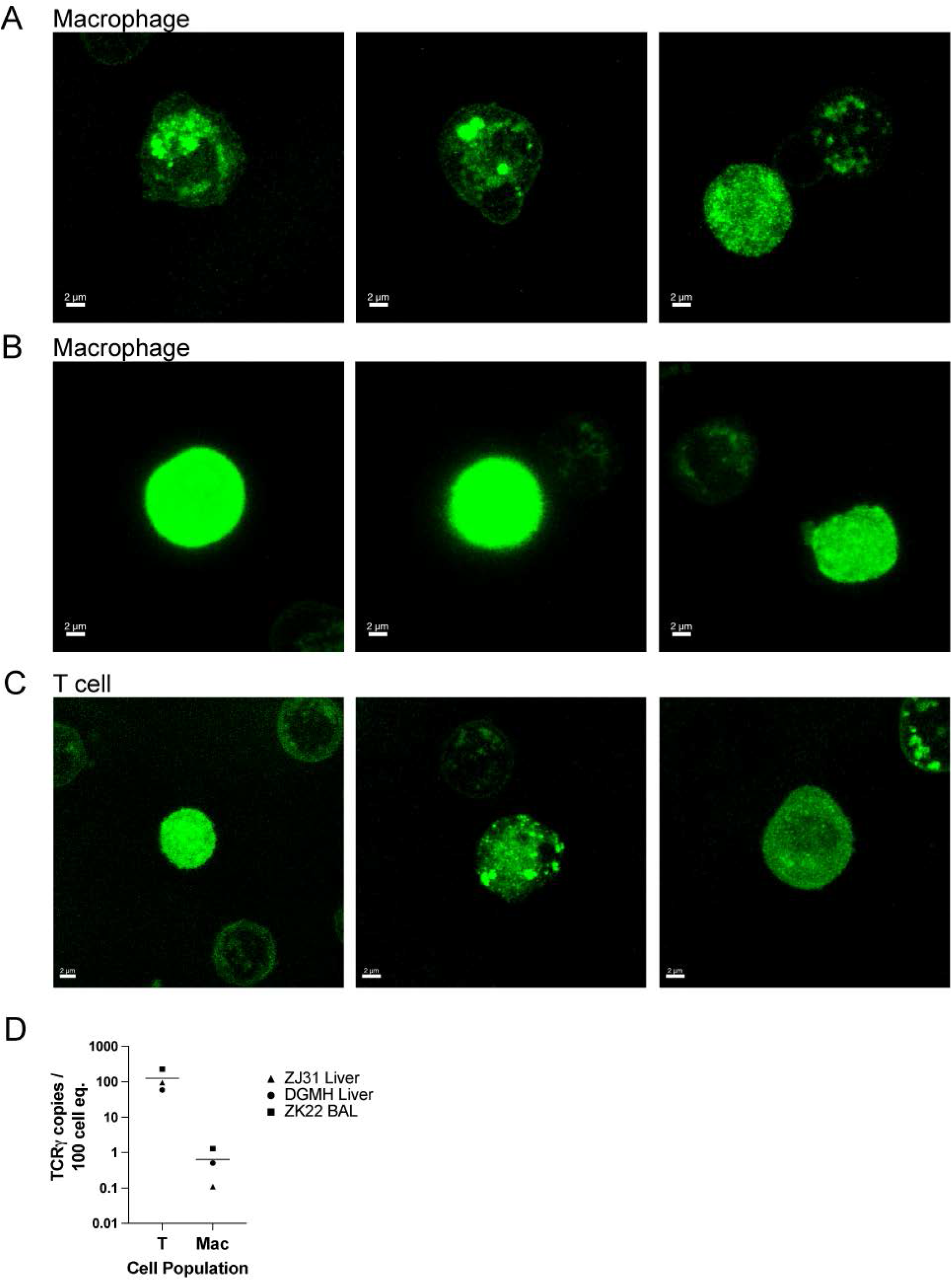
Confocal imaging of GFP localization and analysis of TCR rearrangement within tissue-resident cells excludes phagocytosis as major source of GFP in tissue macrophages. Confocal microscopic images of cellular GFP protein localization in sorted GFP^+^ tissue macrophages from transplanted animal liver and BAL samples demonstrates both (**A**) cellular organelle organization and (**B**) cytosolic GFP expression. (**C**) confocal microscopic analysis of GFP expression patterns in T cells from the same samples are both localized to organelles and/or widespread throughout the cytoplasm. **(D)** Fraction of rearranged T cell receptor DNA in purified T cells and macrophages from liver and BAL of two transplanted (ZJ31, ZK22) and one non-transplanted (DGMH) animals.

### Clonal derivation of tissue-resident macrophages and T cells

We next sought to examine the clonal ontogeny of the GFP^+^ tissue-resident macrophages. We analyzed the clonal relationships of tissue-resident leukocytes of various lineages from the same anatomic sites to each other and to peripheral blood leukocyte populations from the same time point. We flow cytometrically sorted CD3^+^ T cells, CD14^+^ monocytes, and CD14^+^CD163^+^ macrophages from the peripheral blood, and CD3^+^ T cells and MHC class II^+^CD11b^+^ myeloid cells from tissues, including BAL and biopsies from the jejunum, colon, liver and spleen (representative flow cytometric gating shown in Figure 1B) at two to four separate time points from each of four lentivirally-barcoded animals (Table 1).

Barcode retrieval and quantitation was performed as described following PCR amplification, next generation sequencing, and application of custom Python and R pipelines to identify the fractional contributions of individual transplanted HSPC clones to each sorted cellular sample (Espinoza *et al*., 2021; Koelle *et al*., 2017). This approach has been demonstrated to accurately quantitate individual barcode contributions to even highly polyclonal differentiated cell samples, utilizing very high diversity barcoded lentiviral libraries and a targeted transduction efficiency to ensure that >95% of GFP^+^ cells contain a unique barcode, each representing the output from a single transplanted HSPC. We previously reported that following several months of contributions from lineage-restricted short-term engrafting HSPCs, circulating neutrophils, monocytes, and B cells share very similar and stable clonal contributions arising from multipotent long-term repopulating HSPCs (LT-HSPCs) for many years following transplantation (Koelle *et al*., 2017). T cell clonal patterns consist of some contributions from LT-HSPCs matching contributions to other lineages, but also later waxing and waning oligoclonal expansions found only in T cells, representing peripheral memory homotypic responses (Koelle *et al*., 2017).

We performed principal coordinate analysis (PCA) to determine the relatedness of the clonal output contributing to T cells, tissue-resident macrophages and peripheral blood monocytes, and the stability of contributions over several sampling time points. The first principal coordinate separated T cells from myeloid cells and clonal contributions to tissue-resident macrophages were very similar to contemporaneous blood monocytes as shown by clustering on PCA plots (Figure 3A-D). Notably, there was little clonal change over time, at least for the 2-10 months elapsing between sample collection in each animal, and similar clonal patterns between macrophages collected from different anatomic sites. Jejunum and colon samples from animal ZJ31 were less related, however input DNA from low numbers of cells from these tissues likely introduced sampling artifacts due to insufficient cell numbers analyzed to reflect the entire clonal repertoire. T cells from blood clustered separately, as expected, as did T cells from various tissue sites, likely due to responses to local environmental stimuli.

**Figure 3.**
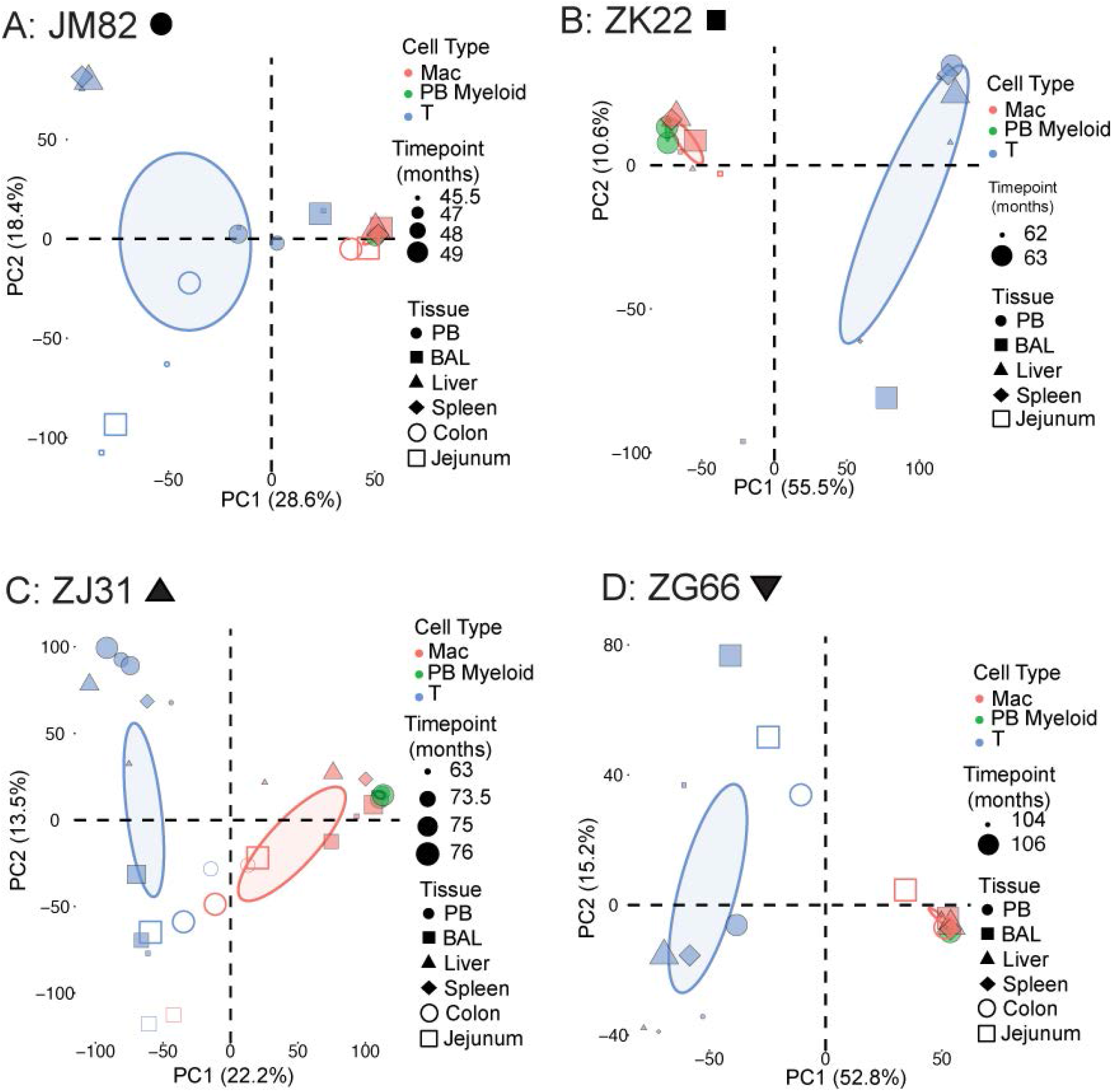
Clonal relationships between tissue-resident cell lineages and peripheral blood monocytes in HSPC-barcoded rhesus macaques. **(A-D)** Principal component analysis of the relationship of the level of all individual barcode reads retrieved, normalized to the total numbers of reads, from tissue-resident macrophages (red), tissue-resident T cells (blue), and peripheral blood monocytes (green) in each animal. Symbol size corresponds to sampling time point relative to transplantation and symbol shape to the tissue type. Ellipses correspond to 95% confidence intervals.

We also performed pairwise Pearson correlation analyses between clonal barcode contributions to blood monocytes, blood macrophages, T cells, and tissue-resident macrophages from various anatomic sites. As shown in Figure 4A-D these correlation analyses confirmed the PCA analyses documenting very highly significant clonal correlations between PB monocytes or macrophages and tissue-resident macrophages, as well as between different anatomic sites and over several time points, and less correlation with circulating or tissue-resident T cells. These data suggest that tissue-resident macrophages develop along the same lineage ontogeny as peripheral blood and circulating monocytes via differentiation from the same set of stable LT-HSPC clones. The similarities in clonal composition between different anatomic sites argues against initial seeding with a small number of precursor cells and ongoing local proliferation/differentiation (Wu et al., 2018a). The low correlation between tissue-resident macrophage and T cell clones also rules out phagocytosis of T cells as an explanation for GFP^+^ of tissue-resident macrophages.

**Figure 4.**
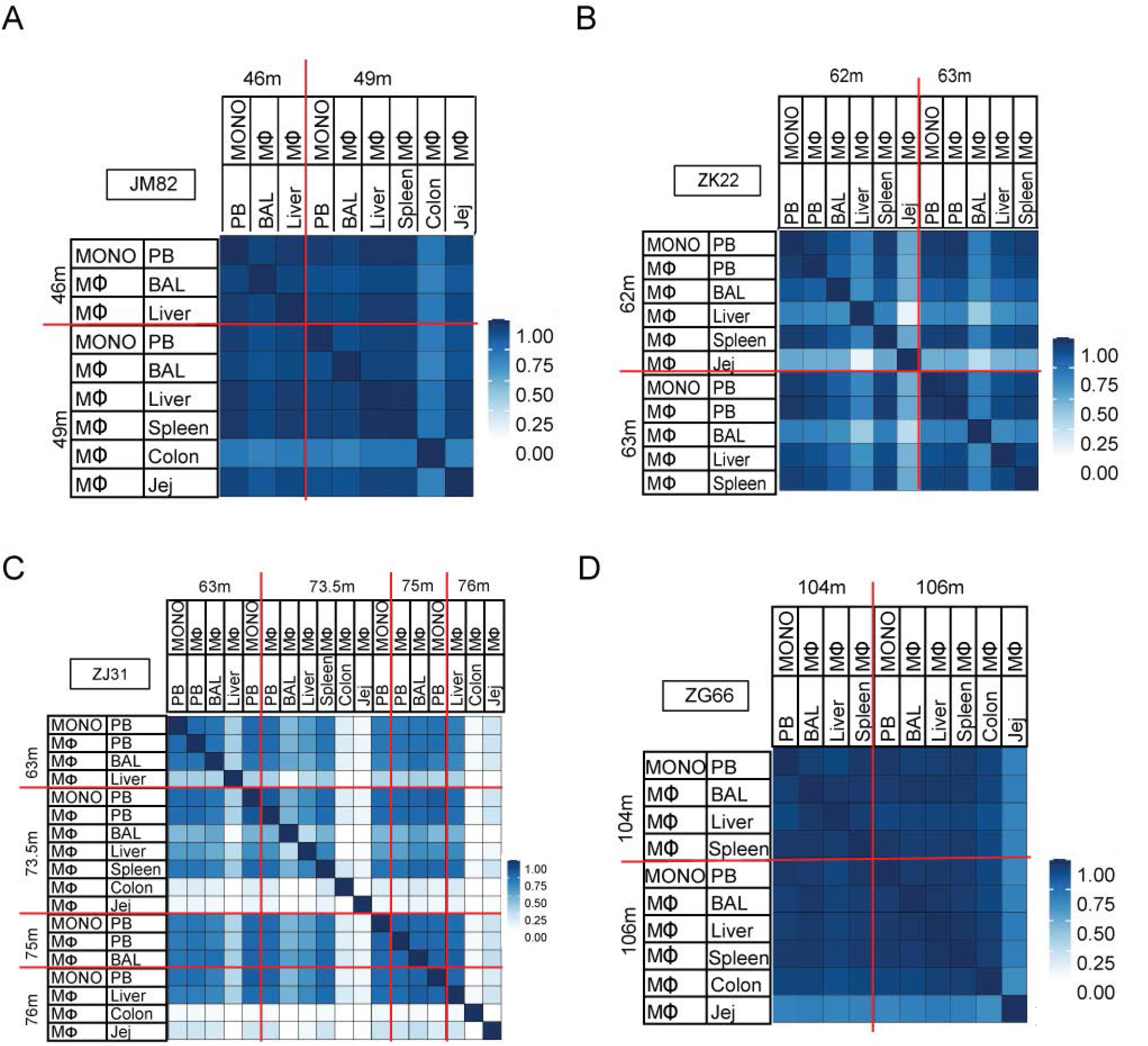
Pairwise comparison of clonality between blood and tissue leukocytes. **(A-D)** Pearson correlation coefficients comparing pairwise fractional contributions between macrophages, monocytes, and T cells in BAL, liver, GI tract, and peripheral blood. The color scale for r values is shown on the right.

To gain insights into the issue of whether seeding of tissues with macrophage precursors such as monocytes occurs only immediately following ablative conditioning with radiation, or occurs on an ongoing basis, we studied clonal patterns over time. While barcodes detected in tissue-resident macrophages and T cells were fairly consistent between the time points sampled (Figures 3-4), these timepoints were 46 months to 106 months following transplantation in the 4 barcoded monkeys. As noted above, we had previously observed and reported that short-lived lineage-biased HSPCs contribute to blood lineages for the first several months following transplantation and then replaced by contributions from stable multipotent LT-HSPCs (Koelle *et al*., 2017; Wu *et al*., 2014). This pattern allowed us to ask whether the HSPC-derived tissue-resident macrophages were progeny of short-term HSPCs cells seeding tissue immediately following ablative conditioning due a unique depletion of tissue-resident macrophages at that time, or were derived instead from LT-HSPC progeny with ongoing replacement over time. Thus, we compared the clonal composition of peripheral blood monocytes sampled for the first 1-2 months following transplantation to those identified from tissue-resident macrophages years following transplantation for each animal (representative longitudinal bar codes isolated from monocytes and tissue macrophages in Figure S1). We performed PCA analysis to determine the degree to which the clones of HSPCs from which T cells, monocytes, and macrophages were related (Figure 5). From this analysis, it was clear that the clones contributing to peripheral blood monocytes and T cells early after transplantation were significantly different from the clones contributing to tissue-resident macrophages and tissue-resident T cells at later timepoints (Figure 5A-D). Again, the first principal coordinate separated T cells from myeloid cells (Figure 5A-D). Moreover, the clones contributing to tissue-resident macrophages most closely resembled the LT-HSPC clones that contributed to development of contemporaneously sampled peripheral blood monocytes as well as other circulating lineages for each animal (Figure 6). Thus, while conditioning TBI may have facilitated initial tissue-resident macrophage turnover and replacement, our model suggests ongoing replacement of these cells from HSPCs.

**Figure 5.**
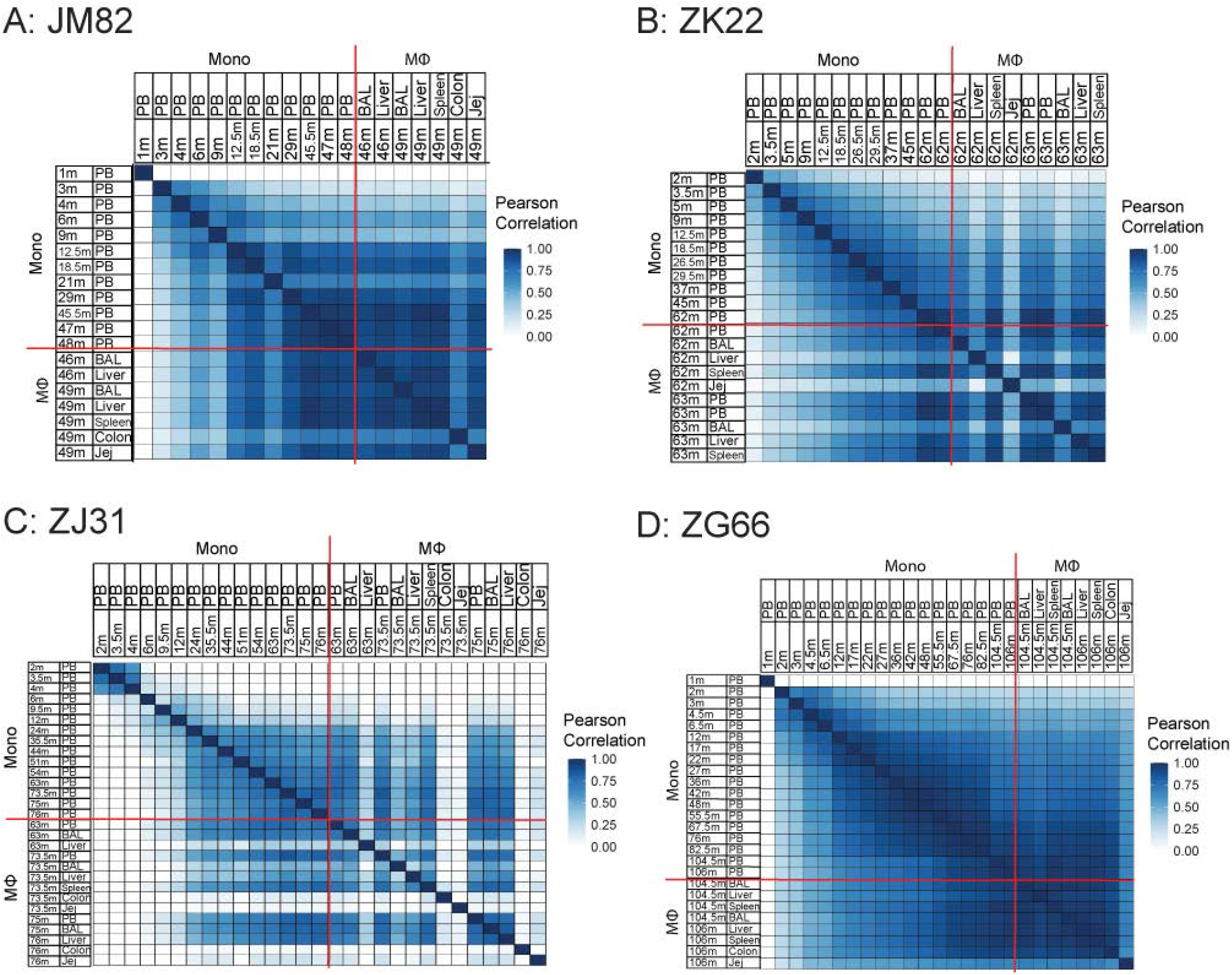
Continual development of tissue-resident macrophages and T cells from HSPCs. **(A-D)** Principal component analysis of the relationship of the level of all individual barcode reads retrieved, normalized to the total numbers of reads, from tissue-resident macrophages (red), peripheral blood monocytes (green), and T cells (blue) from the barcoded animals at multiple timepoints post transplantation. Symbol size represents biopsy time points relative to transplantation, symbol shapes to different tissues, and ellipses correspond to 95% confidence intervals.

**Figure 6.**
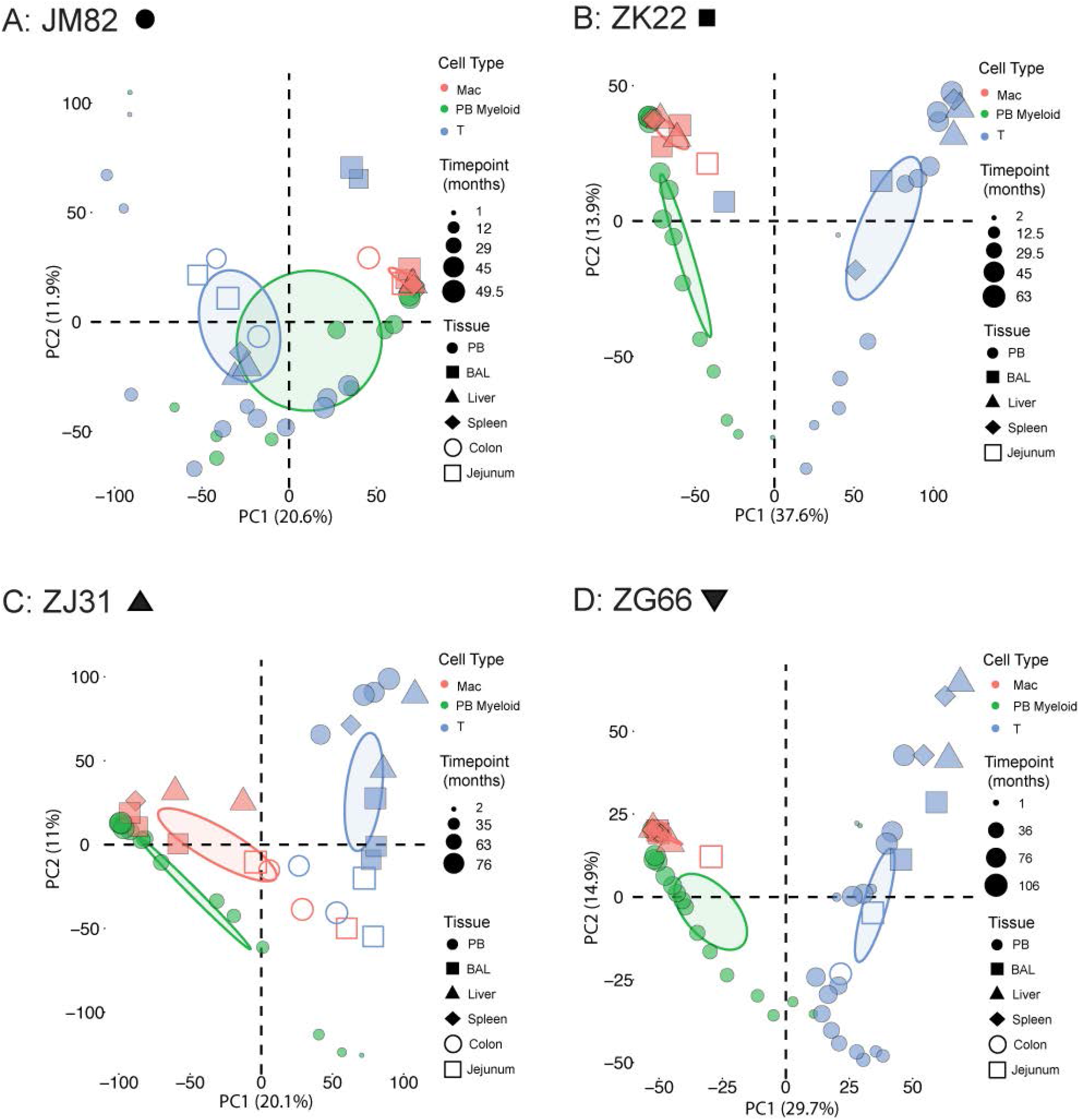
Continual development of tissue-resident macrophages from HSPCs. **(A-D)** Pearson correlation coefficients comparing pairwise fractional barcoded clonal contributions between tissue-resident macrophages and peripheral blood monocytes at multiple timepoints post-transplantation.

### Turnover of tissue-resident macrophages at steady state

Our data clearly demonstrate robust replacement of tissue-resident macrophages after TBI conditioning and autologous transplantation with genetically-modified HSPCs, with ongoing tissue migration and differentiation into tissue-resident macrophages as late as several months after pre-transplantation conditioning, given the shared clonal ontogeny of tissue-resident macrophages with multi-potent long-term HSPCs rather than short-term HSPCs responsible for the initial several months of hematopoietic reconstitution. However, the initial impact of the conditioning regimen could confer conditions longer term that might contribute to increased tissue myeloid cell turnover (Elmore *et al*., 2014; Stutchfield *et al*., 2015; Theurl *et al*., 2016). To investigate the degree of tissue-resident macrophage turnover at steady state, independent of the impact of TBI, we intravenously infused 4 healthy young adult, non-transplanted rhesus macaques with 30 mg/kg/day bromodeoxyuracil (BrdU). On day 10 of infusion, we sampled spleen, and both peripheral and mesenteric lymph nodes. PB myeloid cells and tissue-resident macrophages were analyzed by flow cytometry for incorporation of BrdU into the genome and for active cycling via Ki67 staining (Figure 7). To determine if BrdU was lost from these cells either by cell death or cycles of ongoing proliferation, we sampled the same anatomic sites 24 days following initiation of BrdU (eg 14 days after BrdU infusion was discontinued) (Figure 7). BrdU incorporation was easily detectable in an appreciable percentage of myeloid cells in the blood (41.4 ± 15.7% at day 7, 11.2 ± 4.3% at day 10) and importantly in tissue-resident macrophages at day 10 (31.7 ± 16.4%) but decreased to significantly lower levels by day 24 (6.4 ± 5.1%) (Figure 7). While this analysis could not differentiate between ongoing input from HSPCs or local proliferation of tissue macrophages, these data demonstrate the relatively rapid turnover of lymphoid tissue-resident macrophages at steady state and support the concept that continuous replacement of these tissue-resident macrophages is ongoing, even in the absence of conditioning.

**Figure 7.**
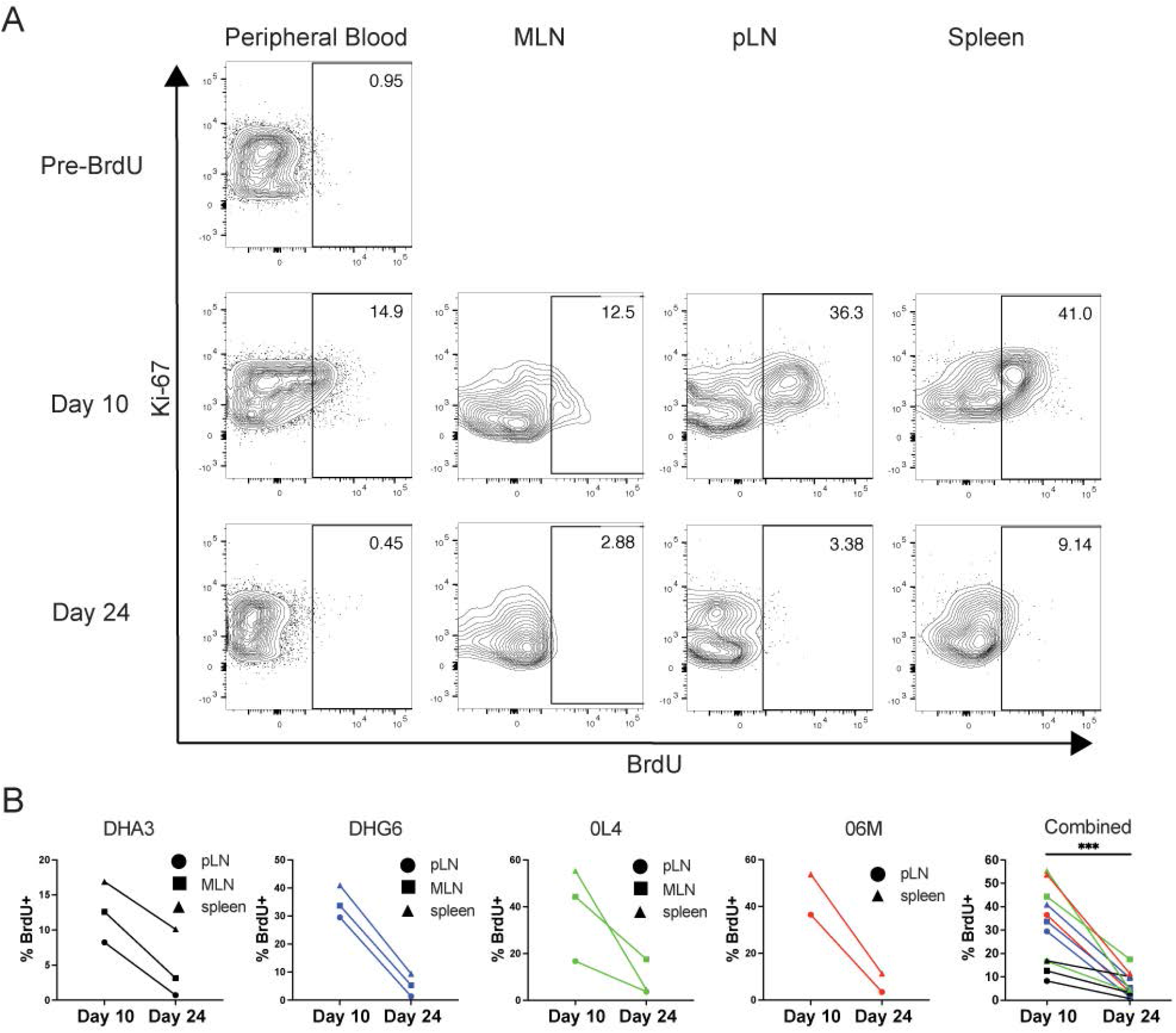
Tissue macrophages turnover at steady state. Healthy, non-transplanted, rhesus macaques were infused with bromodeoxyuracil (BrdU) intravenously for 5 days. Five days from the conclusion of administration (Day 10) and 14 days later (Day 24), blood and tissue cells were analyzed by flow cytometry to assess BrdU incorporation. (**A)** Representative plots showing percent BrdU positive of at least 1000 cells defined as single, live, CD45^+^, CD11b^+^, HLA-DR^+^, CD20^-^ mononuclear from the same animal at each timepoint. (**B)** BrdU levels from all four healthy non-transplanted animals in tissue macrophages. *** p <0.001, Wilcoxon matched-pairs test.

## Discussion

Macrophages regulate tissue immunity, orchestrating the initiation and resolution of antimicrobial immune response, as well as the maintenance of tissue integrity. Their ontogeny and lifespan have been the subject of intense interest given their broad range of functions and phenotypes in health and disease (Ortiz et al., 2015), including regulation of tissue repair (Wynn and Vannella, 2016), susceptibility to infection by DNA and RNA viruses (Estes *et al*., 2018), control of local tissue immune responses with an ability to retain epigenetic “memory” of prior infection (Bekkering et al., 2021), transformation to an immunosuppressive role by tumors (Hourani et al., 2021), and neoplastic transformation (Varol et al., 2015). Such understanding is necessary to intervene in these processes for clinical benefit.

Studies in mice have revealed that embryonically macrophages are first directly generated in the yolk sac without monocytic precursors. Murine models which allow for fate mapping and the ability to selectively delete genes at different stages of differentiation (and in particular cell types) has been invaluable to our understanding of immunological development. Upon establishment of the blood circulation, yolk sac macrophages seed the whole embryo (McGrath et al., 2003). A paradigm emerged suggesting that these embryonically-derived macrophages persist into adulthood and continue to constitute the bulk of the adult tissue macrophage compartment (Hoeffel and Ginhoux, 2018). Murine models with specific impairment of definitive but not primitive hematopoiesis demonstrated that yolk sac–derived cells can indeed contribute to tissue-resident macrophage populations in adult mice (Schulz *et al*., 2012). Parabiosis and some depletion studies suggested that once seeded with embryonic macrophages, tissue-resident populations are maintained by local proliferation, not via contributions from circulating monocytes (Hashimoto et al., 2013). However, some models instead suggest that murine tissue-resident macrophages in adult mice require ongoing replenishment from HSPCs (Sheng et al., 2015) and the topic remains controversial.

While murine studies have driven our fundamental understanding of immunological processes and mammalian development and mechanisms underlying multiple diseases, there are accumulating data suggesting that highly inbred and caged mice housed under SPF conditions can fail to capture important genetic and environmental factors that influence physiology (Masopust et al., 2017), including tissue-resident macrophage biology. SPF mice reconstituted with the GI tract microbiome of wild mice can survive influenza challenges that are lethal in SPF mice (Rosshart *et al*., 2017). Tissue-resident macrophages in SPF mice have been shown to die and then rapidly repopulate under stress conditions. Kupffer tissue resident macrophages in the liver recycle iron from dying red blood cells (RBC), and intravenous administration of a large load of dying RBCs results in iron-related toxic death and subsequent repopulation of almost every Kupffer cell (Theurl *et al*., 2016). Aspirin toxicity similarly leads to significant turnover of Kupffer cells, dependent on expression of the colony stimulating factor 1 (CSF1) receptor (Stutchfield *et al*., 2015). Similarly, CSF1R signaling is critically important for the maintenance of microglia cells in the brain, and blockade of CSF1R lead to loss of >99% of microglia cells, with rapid replenishment from nestin^+^ cells (Elmore *et al*., 2014). Finally, administration of chemotherapeutic agents, aimed at killing rapidly-dividing neoplastic cells but also leading to the death of multiple haematopoietically-derived cells that proliferate *in vivo*, also results in loss of microglia cells in the brain which are subsequently replenished from peripheral myeloid cells (Sailor *et al*., 2022). Taken together, it is clear that even in mice, macrophages can be short lived and rapidly replenished from monocyte precursors in peripheral blood in certain settings.

Given the drawbacks of extrapolating murine results to human biology and disease, it is of great importance to consider these questions in humans and in outbred non-human primate models animal models. To date, there have been few studies of the ontogeny of tissue macrophages in adult humans or primates. In the context of allogeneic HSPC transplantation, a few early studies used low resolution approaches in sex-mismatched donor-recipient pairs to demonstrate donor-type perivascular myeloid cells in the brain (Unger et al., 1993). However, graft-versus host ablation of host-type HSPC but also mature hematopoietic cells makes interpretation of allogenic tissue-resident macrophage chimerism difficult to extrapolate to a non-allogeneic transplantation setting. However, the efficacy of allogeneic transplantation and of autologous HSPC gene therapy in the treatment of some central nervous system metabolic storage disorders such as adenoleukodystrophy suggest that replacement of at least the function of microglial cells and other tissue resident macrophages by HSPC-derived cells is possible (Eichler et al., 2017; Miller et al., 2011). An early retroviral gene marking study in RMs demonstrated GFP-expressing perivascular macrophages in the brain (Soulas et al., 2009).

Our current study provides detailed and unequivocal evidence that significant percentages (up to 50%) of macrophages resident in lung (BAL), liver, GI tract, and lymphoid tissues can be derived from HSPC precursors in adult NHPs. The clonal tracking results in our model strongly support the concept of continuous replacement of macrophages by circulating long-term HSPC-derived cells. Even in the absence of transplantation, the labeling studies with BrdU demonstrate ongoing rapid turnover of tissue-resident macrophages at steady state. Importantly, given that the transduction efficiency of HSPCs is less than 100% (Wu *et al*., 2018a; Wu *et al*., 2014), our analysis underestimates the total number of tissue-resident macrophages that had developed from HSPCs.

These data are consistent with mathematical modelling of decay rates of virus production from macrophage-tropic HIV/SIV infections of immunocompromised mice transplanted with a human immune system and of nonhuman primates (Honeycutt et al., 2017; Micci et al., 2014). In these studies, it was demonstrated that virus-infected macrophages had an incredibly short half-life, as little as one day. In sum, these results of great relevance to consideration of how to address HIV reservoirs in infected humans.

It remains possible that some of the tissue-resident, GFP^-^, macrophages in our transplanted NHPs contained cells were extremely long lived and derived from non-adult HSPC and represent residual embryonic-macrophage derived cells. Moreover, our analysis was, by no means, extensive of all anatomic sites in which macrophages reside. Thus, future studies are certainly warranted, including ongoing analyses of brain microglial populations, a topic of intense current interest. Irrespective, our results demonstrate that HSPCs continuously develop into tissue-resident macrophages in adult NHPs, that these cells turnover at steady state, and the feasibility of such analyses in primates. Data from these studies will aid in development of therapeutic interventions aimed at attenuating the function and lifespan of tissue-resident macrophages in health and disease.

## Author Contributions

ARR, CW, SH, LP, HDH, and JMB performed experiments for the manuscript. ARR, CW, LP, and HDH analyzed the data and created figures. JMB and CED conceived and supervised the study. All authors contributed to the writing and editing of the manuscript.

## Acknowledgements

Funding for this study was provided by the Divisions of Intramural Research of the National Institute of Allergy and Infectious Diseases and the National Heart, Lung, and Blood Institute. The content of this publication does not necessarily reflect the views or policies of the U.S. Government, nor does the mention of trade names, commercial products, or organizations imply endorsement by the U.S. Government. The authors would like to thank the NHLBI, NIAID, and DVR non-human primate facility veterinary staff for skilled animal care and tissue sampling, and the NHLBI DNA sequencing and genomics and flow cytometry cores.

## Notes

### Competing Interest Statement

The authors have declared no competing interest.

