## Supplemental Table 1 for "Ongoing Production of Tissue-Resident Macrophages from Hematopoietic Stem Cells in Healthy Adult Macaques"

**Supplemental Table S1. Reagents used for flow cytometric analysis**

| **Antibody** | **Fluorochromes** | **Clone** | **Company** |
|---|---|---|---|
| Anti-CD3 | Alx700, BV786, V400 | SP34-2 | BD |
| Anti-NKG2a | APC, PE-Cy7 | Z199 | Beckman Coulter |
| Anti-CD45 | V450 | DO58-1283 | BD |
| Anti-CD11b | BV785 | ICRF44 | BD |
| Anti-CD20 | BUV395 | 2H7 | BD |
| Anti-CD20 | APC-Cy7 | L27 | BD |
| Anti-HLADR | APC-H7 | G46-6 | BD |
| Anti-CD49d | PE | TS2/7 | Biolegend |
| Anti-CD16 | BV605 | 3G8 | Biolegend |
| Anti-CD14 | PE, Pacific Blue | TuK4 | Invitrogen |
| Anti-CD163 | BV421 | GH1/61 | Biolegend |
| Live Dead exclusion | Aqua Blue | N/A | Invitrogen |
