## Supplementary figures and images for "Ongoing Production of Tissue-Resident Macrophages from Hematopoietic Stem Cells in Healthy Adult Macaques"

### Supplemental figure 1

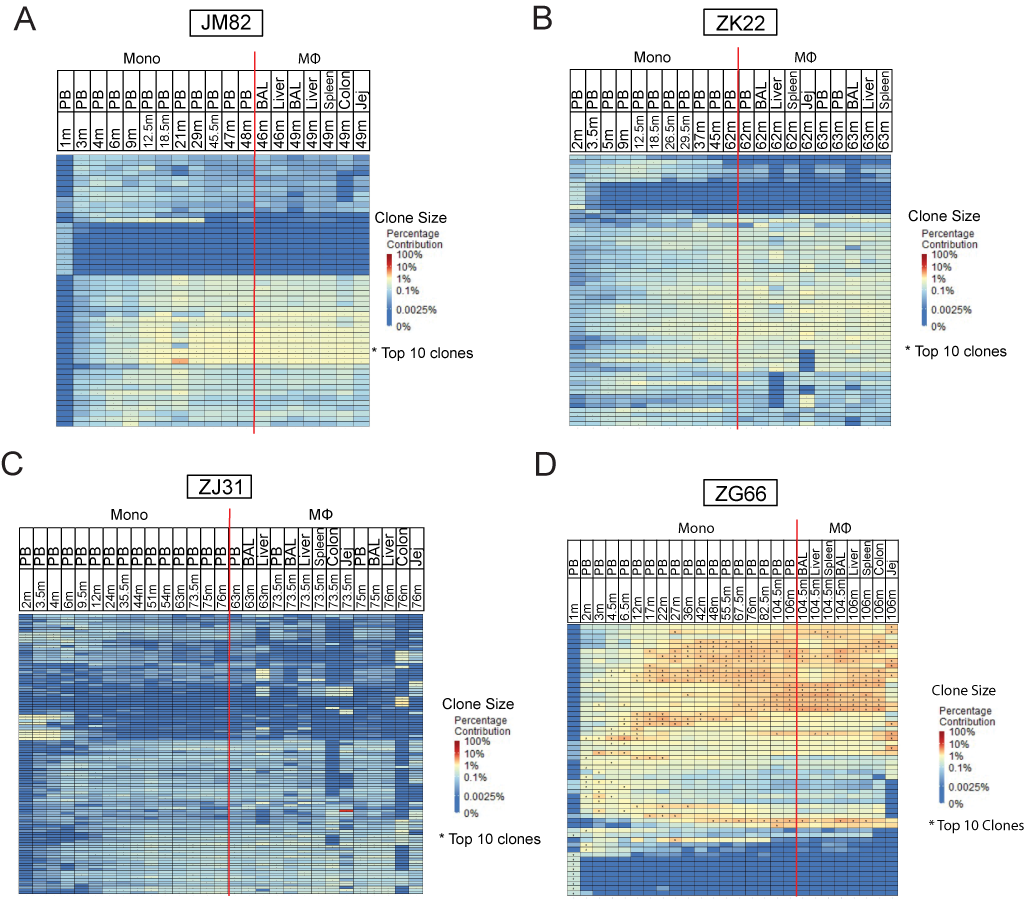
